# Understanding the physiological alterations of *Vibrio cholerae* upon exposure to L-ascorbic acid

**DOI:** 10.64898/2026.08.31.748307

**Authors:** Manpreet Kaur, Sakshi Gautam, Suman Paul, Himanshu Sen, Surajit Bhattacharjee, Saumya Raychaudhuri

## Abstract

The scourge of cholera remains a major global public health threat. It affects up to 4 million people worldwide and causes tens of thousands of deaths each year. The disease is experiencing a concerning resurgence in many parts of Africa, the Middle East, and Asia. To effectively tackle cholera and circumvent rising antimicrobial resistance, targeted biological and preventive approaches, complementing traditional rehydration, are urgently needed. In this regard, our group has demonstrated the efficacy of L-ascorbic acid in controlling the growth and pathogenesis of *Vibrio cholerae in vitro*. The present work further provides a mechanistic elucidation of the L-ascorbic acid-mediated physiological changes in *V. cholerae* and also bolsters such a non-antibiotic approach to control cholera.

## Introduction

*Vibrio cholerae* is the pathogen responsible for cholera, a disease found worldwide. Every year, millions of people across the globe contract this deadly illness [1]. The disease significantly affects the socio-economic status of affected nations, especially in poverty-stricken and war-affected areas [2–4]. A recent study on the global mortality rate shows an upward trend [5]. Food security, rather than food and nutrition insecurity, is one of the social determinants of physical and mental health. Intriguingly, food insecurity is also linked with an increased risk of cholera [6]. These studies collectively emphasise the need to develop more effective measures to prevent the disease.

Decades of research on *V. cholerae* reveal numerous remarkable facts about the organism’s biology, particularly concerning its epidemiology, pathogenesis, and survival in various hosts and aquatic environments [7]. Among the 220 serogroups of *V. cholerae*, strains belonging to O1 and O139 are associated with epidemics globally. The other remaining serogroups, termed non-O1/non-O139, are generally associated with cases of sporadic diarrhoeal infections [8,9]. Marked by severe dehydration, the primary treatment for cholera has traditionally involved rehydration therapy, either through oral rehydration solution (ORS) or intravenous fluid supplementation. This forms the mainstay of the cholera treatment regimen. In addition to ORS, antibiotics are also necessary as adjunct therapy to control and reduce the duration of the outbreak. The use of antibiotics also increases the incidence of drug-resistant *Vibrio cholerae* strains [10,11]. Currently, three WHO-approved oral cholera vaccines, Dukaral, Shankol, and Evichol, are available in the market. However, these vaccines require two doses to achieve full protection, and none of these OCVs is recommended for infants. The present focus is on developing a single-dose immunization with an improved formulation to protect young children [12,13]. Therefore, there is a demand for newer and more effective strategies to tackle this deadly disease.

Previously, we showed the effectiveness of L-ascorbic acid (L-AA) in restricting the growth of *V. cholerae* under *in vitro* and *in vivo* experimental conditions [14,15]. In this study, we aimed to investigate the effect of stress originating from L-AA on the overall physiology of *V. cholerae*. We documented L-AA-mediated production of reactive oxygen species (ROS) in our assay. Microscopically, we observed significant changes in the membrane architecture of *V. cholerae*, which further led to an increase in membrane permeability. Global transcriptomic analysis also revealed differential regulation of several genes in response to this stressor. Collectively, exposure to L-ascorbic acid caused significant physiological changes in *V. cholerae*, thus affecting the growth of the organism.

## Material and Methods

### Bacterial strain and media

This study was conducted on the *Vibrio cholerae* strain N16961 O1, El Tor, Ogawa, which was generously provided by Andrew Camilli, Tufts University. The strain was maintained in Luria-Bertani broth (LB) (Difco) supplemented with 0.1 mg/mL streptomycin under shaking conditions at 37°C, 200 rpm or on LB agar at room temperature.

### Estimation of Reactive Oxygen Species (ROS)

Production of intracellular ROS in *V. cholerae* after treatment with L-AA (Sigma-Aldrich) was estimated through 2’,7’-dichlorofluorescin diacetate (DCFDA) fluorescence dye following the published protocol of [16]. Log-phase *V. cholerae* culture was diluted and treated with L-AA 2.5 mg/mL and 5 mg/mL and incubated for 1 h at 37□. Following incubation, cells were centrifuged and washed with 1X PBS and resuspended again in the 1X PBS and added DCFDA at a concentration of 10 µM and incubated in the dark for 30 min at room temperature. The fluorescence intensity was measured at excitation 484 nm and emission 525 nm using a BioTek Synergy H1 Hybrid microplate reader [17].

### Scanning electron microscopy analysis

The impact of L-AA on cell morphology was examined using field emission scanning microscopy (FESEM) following the method described previously with minor changes [18,19]. The overnight-grown *V. cholerae* N16961 was inoculated in LB medium with appropriate antibiotics to attain the mid-log phase (OD_600_ = 0.4–0.6). Cells were collected by centrifugation at 5000 rpm for 10 min, followed by washing twice with 0.1 M PBS, and then resuspended in LB medium containing L-AA at final concentrations of 2.5 mg/mL and 5 mg/mL. The cultures were incubated at 37°C, 200 rpm for 1 h. After the stipulated time, cells were pelleted, washed twice with 0.1 M PBS, and fixed with 4% glutaraldehyde (Sigma Aldrich) at 4°C for 12 h. After fixation, cells were gently washed three times using 0.1 M PBS and subjected to graded ethanol dehydration steps (30%, 50%, 60%, 70%, 80%, 90%, and 100%). Finally, cells were reconstituted in 100% ethanol, and the suspension was spread on the glass slide. Samples were air-dried in a desiccator overnight at room temperature. The fixed slides of all the samples were examined using an Apreo S FESEM (Thermo Scientific, USA). A small amount of gold was coated on the samples to prevent charging in the microscope. Secondary electron images were captured at low energies between 2 keV and 2.5 keV.

### Transmission electron microscopy

Similarly, after treatment, samples were collected, washed three times, and mixed in 0.1 M PBS. For microscopy, the samples were loaded on the carbon-coated copper grid (TED PELLA) for 10 min at room temperature. The excess sample was removed by filter paper. After that, the negative staining was performed with 2% uranyl acetate (Sigma-Aldrich) for 30 s, followed by removal of excess stain and putting the grid in the air for drying. The samples were viewed under JEOL-TEM 2100 @ 200 kV HT.

### Effect of L-ascorbic acid on the cell membrane

Membrane permeabilities were monitored using propidium iodide (PI) staining as previously described, with slight modifications [20,21]. Following treatment, cells were obtained by centrifugation at 5000 rpm for 10 min. Cells were then washed twice with 1X PBS and mixed in the same buffer to an OD_600_ of 0.3. Propidium iodide (PI) was added to 200 μL of the cell suspension to obtain the final concentration of 5 μg/mL, and the cell suspension was transferred to a black 96-well microplate (Greiner Bio-One, MICROLON®). The cells were then placed in the dark for 10 min, and PI fluorescence intensity was measured using the Synergy Hybrid plate reader (BioTek) at excitation/emission wavelengths 580/620 nm.

### Total RNA purification and RNA sequencing

The overnight cultures of *V. cholerae* N16961 were taken to set up the secondary cultures in LB media with appropriate antibiotics and grown till log phase (OD□ □ □ = 0.4-0.6). Culture was then pelleted down and washed twice with 1X PBS, resuspended in LB media and set up the experiment at O.D. 600 = 1.0 with untreated and treated N16961 with 2.5 mg/mL L-AA. Incubate the cultures at 37°C, 200 rpm for one hour. The samples were pelleted down, and total RNA was isolated using the RNeasy mini kit (Qiagen Technologies, Hilden, Germany) according to the manufacturer’s guidelines with minor modifications [22]. After the purification, RNA concentrations were measured on a Nano Drop (Thermo-fisher, USA).

The samples were processed for quality control assessment, library preparation, sequencing, annotation and data analysis were carried out by Strand Life Sciences (Bangalore, India). Initially, libraries were generated using the Kapa Hyper Plus kit, and sequencing was performed using NovaSeq X Plus to generate 60 million reads per sample with a read length of 150 bp. The generated data can be accessed through the NCBI Sequence Read Archive (SRA) database using reference ID PRJNA1461801.

### Assembly and functional enrichment analyses of differentially expressed genes

The demultiplexed FASTQ files were imported into Strand NGS v4.1 for alignment and further analysis. Briefly, Pre-alignment Quality control QC was performed, followed by trimming of 10 bases at the 5′ end. Reads were mapped to the reference genome and the transcriptome of *V. cholerae* O1 biovar El Tor strain N16961 (assembly GCF_000006745.1, February 2025), and post-alignment QC confirmed mapping reliability. Gene-level counts were generated and normalized using the Trimmed Mean of M-values (TMM) method. Lowly expressed genes were filtered by retaining those with ≥3 reads in at least 2 of 6 samples. Differentially expressed genes (DEGs) analysis was performed using EdgeR to identify genes exhibiting statistically significant expression using the quasi-likelihood (QL) framework. The p-values were corrected using the Benjamini-Hochberg method, with a false discovery rate (FDR) threshold of 0.05. Genes with a fold change (FC) ≥2 were classified as upregulated, whereas those with FC ≤−2 and p-value <0.05 were considered as downregulated.

### Volcano plot and heat map

Volcano plot visualization of DEGs was performed using *Python* (version 3.12.2) with the *pandas* and *matplotlib* libraries. Log2 fold change (log□FC) values were plotted against −log□ □-transformed p-values. Heatmaps were created using Python (*pandas, seaborn, matplotlib*). Log2 fold change values were Z-score normalized using *StandardScaler*, and Hierarchical clustering of genes was performed using *seaborn clustermap*. Samples were annotated based on treatment conditions, and column clustering was disabled to preserve group structure.

### GO and KEGG enrichment analysis

For pathway enrichment analysis, these locus tags were first functionally annotated using eggNOG-mapper to obtain KEGG Orthology (KO) identifiers and Gene Ontology (GO) terms. KO identifiers represent functional orthologs associated with biological pathways, while GO terms describe gene functions across biological, molecular and cellular processes. KEGG pathway and GO pathway enrichment analyses were visualized in R using the *ggplot2, dplyr*, and *stringr* packages. Adjusted p-values were transformed to −log10 values to facilitate visualization of enrichment statistical significance. For KEGG analysis, the top 20 significantly enriched pathways ranked by −log10 (adjusted p-value), were selected for visualization. For GO analysis, the top 15 significantly enriched terms, ranked by adjusted p-values, were selected for visualization, and separate plots were generated for the Biological Process (BP), Molecular Function (MF), and Cellular Component (CC) categories.

### Quantitative reverse transcription-polymerase chain reaction (qRT-PCR)

The total RNA was isolated from the untreated and treated *V. cholerae* N16961 cells by using the RNeasy mini kit (Qiagen Technologies, Hilden, Germany) according to the manufacturer’s guidelines with minor changes. Gene expression analysis was carried out using a superscript III platinum SYBR green mix one-step qRT-PCR kit (Invitrogen, USA). The complementary DNA (cDNA) synthesis and quantitative PCR amplification were performed in a single reaction mixture containing gene-specific primer pairs (200 nM), and a list of primers used is provided in **Table 1**. RT qPCR was performed using a qTOWER^3^ G Real-Time PCR System (Analytik Jena, Germany) under the following cycling conditions: 50°C for 25 min followed by 94°C for 2 min. Amplification was carried out for 40 cycles, consisting of 94°C for 15 s, 55°C for 30 s, and 72°C for 1 min and at the end, the melt curve was used to confirm the specificity of the amplified products. The endogenous control *V. cholerae* 16S rRNA gene was used, and relative fold changes were calculated using the 2^−ΔΔCt method [23].

**Table 1.**
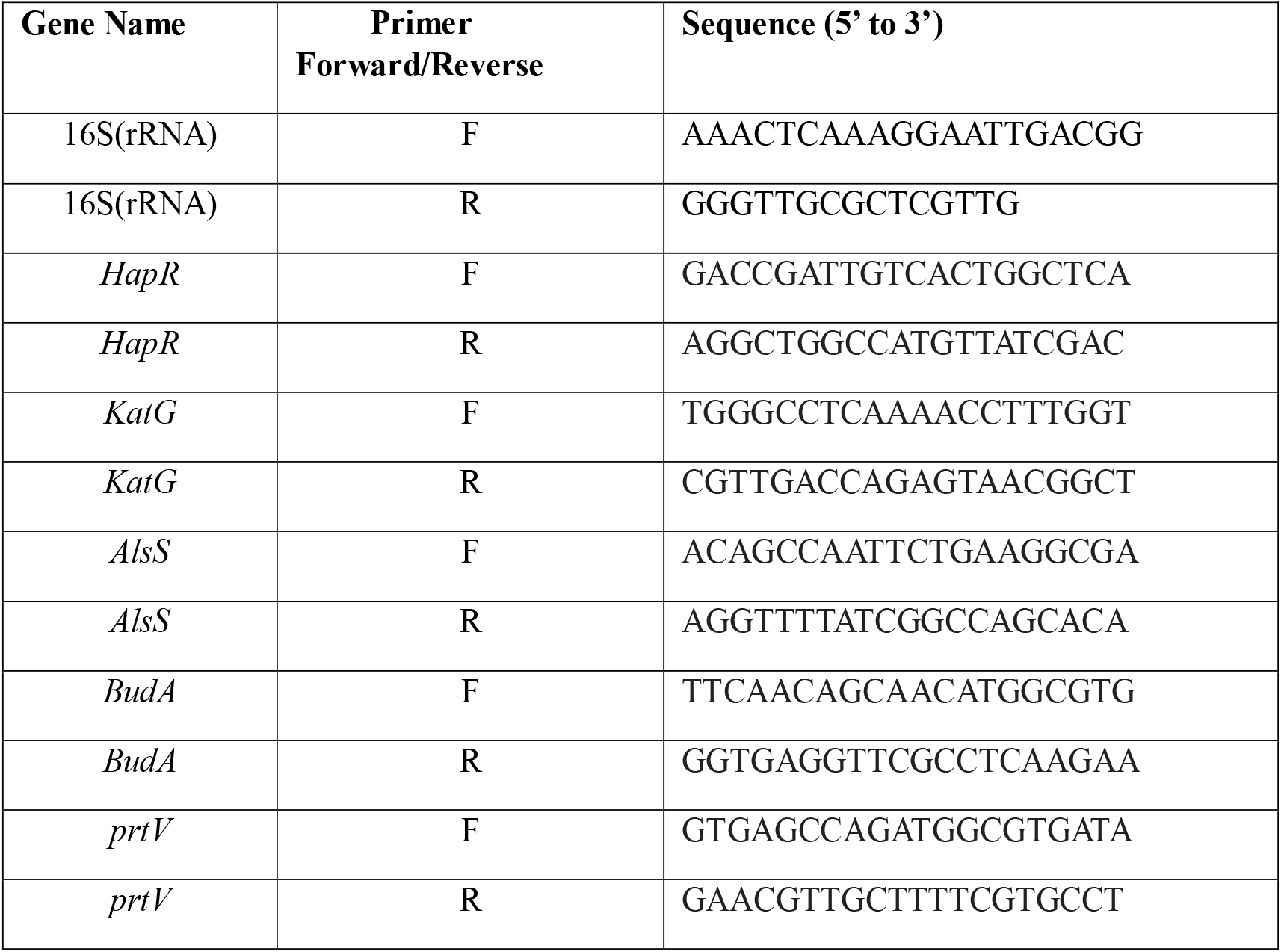
List of primers used for this study.

## Results

### L-ascorbic acid damages the membrane integrity of *V. cholerae*

Previously, we showed the effectiveness of L-ascorbic acid (L-AA) in controlling the growth of various *V. cholerae* strains under different growth conditions [14]. The antimicrobial activity of vitamin C is now well documented [24–26]. It has been demonstrated that L-AA exhibits both antioxidant and prooxidant properties. The prooxidant property appears to contribute to the antimicrobial activity of L-AA, as reported in previous studies [26,27]. To assess ROS production by L-AA under our experimental conditions, we followed the published protocol (Xu et al., 2022) to measure ROS using the fluorescent dye DCFDA. Our data also confirmed that ROS production occurred under our assay conditions (**Figure 1A**). L-AA can disrupt bacterial cell membranes, as evidenced in the case of *Staphylococcus aureus* [28]. To estimate the extent of damage, we performed electron-microscopic analysis of the L-AA-treated *V. cholerae* strain N16961. Both scanning electron microscopy (SEM) and transmission electron microscopy (TEM) images showed normal, rod-shaped, smooth, morphologically intact cell membranes in untreated cells. We observed significant morphological changes in cells after treatment with different concentrations of L-AA. Treatment with low concentrations of L-AA (2.5 mg/mL) resulted in loss of the curved rod shape, cell shortening and distortion, surface roughening, membrane blebbing, cell shrinkage and deformation. At the higher concentration of L-AA (5 mg/mL), severe morphological damage was evident, including membrane disruption, deformation, surface wrinkling and corrugation, loss of structural integrity, and exudation of the contents (**Figure 1B and C**). Overall, these observations underscore that L-AA induces dose-dependent structural damage in *V. cholerae*, leading to loss of cell integrity and morphological disruption.

**Figure 1.**
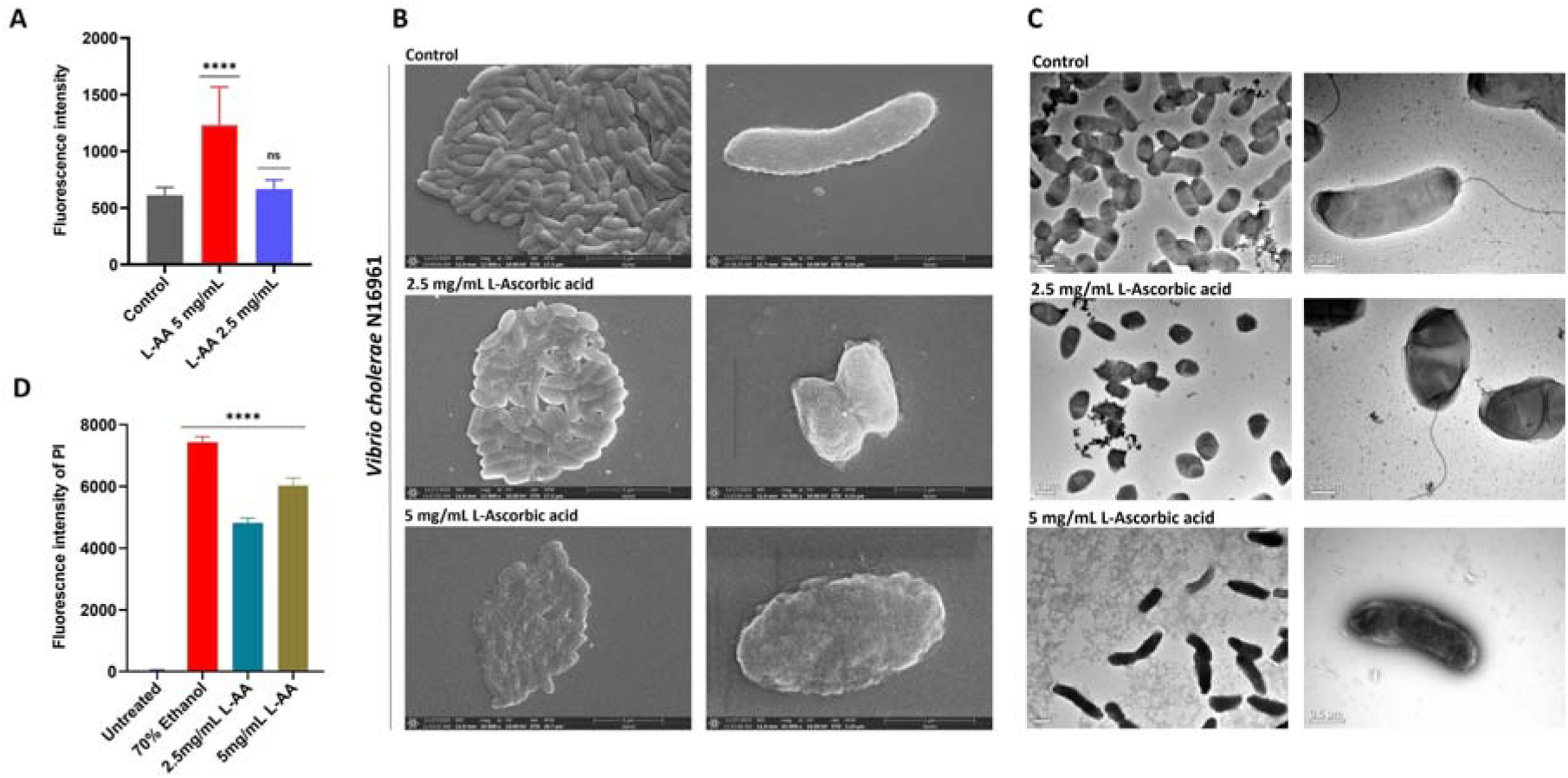
Effect of L-ascorbic acid on oxidative stress, membrane integrity and cell morphology in *Vibrio cholerae* N16961: **(A)** Estimation of intracellular ROS in *V. cholerae* using DCFDA fluorescence dye. Effect of L-ascorbic acid (L-AA) 5 mg/mL and 2.5 mg/mL on the production of intracellular ROS in *V. cholerae*. Data were analysed using one-way ANOVA, and the significance level was indicated as ****p□0.0001 **(B)** Cells of *V. cholerae* N16961, untreated and treated samples with different concentrations of L-AA, i.e., 2.5 mg/mL and 5 mg/mL, were visualised under the scanning electron microscope (SEM), which was operated at 10 kV with scale bars of 1 µm and 5 µm. **(C)** In transmission electron microscopy (TEM) images of untreated and treated cells. Cells were placed on the carbon-coated copper grid and negatively stained with 2% urinyl acetate, followed by air drying. Images were acquired at 200 kV HT with a scale bar of 0.5 and 1 µm. **(D)** The membrane permeability assay was performed using propidium iodide (PI). Untreated *V. cholerae* N16961 cells, cells treated with 70% ethanol (positive control), and cells treated with different concentrations of L-AA were incubated with PI (5 μg/mL) for 10 min. Fluorescence intensity was measured at excitation/emission wavelengths of 580/620 nm. Data were analysed using one-way ANOVA, and significance was indicated as ****p < 0.0001.

It should be noted that damage to the bacterial cell wall increases membrane permeability, often leading to leakage of intracellular components, uncontrolled osmotic influx, and cell death. To evaluate the status of membrane permeability, we performed a propidium iodide uptake assay as described elsewhere [16,20]. Cells with an undamaged cell membrane exhibited minimal or no fluorescence intensity, which indicates the intactness of the cell membrane in untreated cells. On the other hand, cells treated with 70% ethanol (as a positive control) showed an increase in fluorescence intensity and complete membrane disruption. Comparing the cells of untreated and treated with L-AA, a significant increase in PI uptake was observed at both concentrations (2.5 and 5 mg/mL), which suggested marked cell membrane damage in a concentration-dependent manner (**Figure 1D**).

### Transcriptional landscape of *V. cholerae* after exposure to L-ascorbic acid

Acid and oxidative stress trigger significant transcriptomic changes in bacteria, triggering the upregulation of genes involved in stress response and the downregulation of others, including those related to metabolism and growth [29]. For example, L-AA effectively inhibited and disrupted *E. coli* biofilm formation by ROS generation, interfering with quorum-sensing regulation and reducing exopolysaccharide (EPS) production [30]. L-AA-mediated inhibition of the growth and biofilm formation has also been evidenced in the case of multidrug-resistant (MDR) *Burkholderia cepacia* [31]. To gain an insight into the transcriptional changes caused by L-AA in *V. cholerae*, we performed a global RNA expression analysis of the cholera bacterium after exposure to L-AA. The comprehensive transcriptomic analysis was performed to identify the mechanism and underlying pathways that were affected in the cholera bacterium after exposure to L-AA. Differentially expressed genes (DEGs) were evaluated by comparing the culture treated with L-AA 2.5 mg/mL to the untreated (control) group. Genes were considered differentially expressed if they showed significant expression by log2 fold change ≥2 or ≤−2 and p < 0.05. A total of 889 genes were differentially expressed in the treated group, including the 429 genes that were upregulated, mostly associated with oxidative stress (*katG, grxB*), acid stress (*cadB*), and metabolic stress (*budA, alsS)*, efflux mechanism (*emrD*) and DNA repair gene (*recR, ruvA*). In contrast, a total of 460 genes were significantly downregulated, mainly associated with energy metabolism *(odhB, citC*), membrane modification (*almF*), virulence (*vasJ, acfC*) and quorum sensing (*hapR, cqsA, cqsS*) (**Figure 2A**).

**Figure 2.**
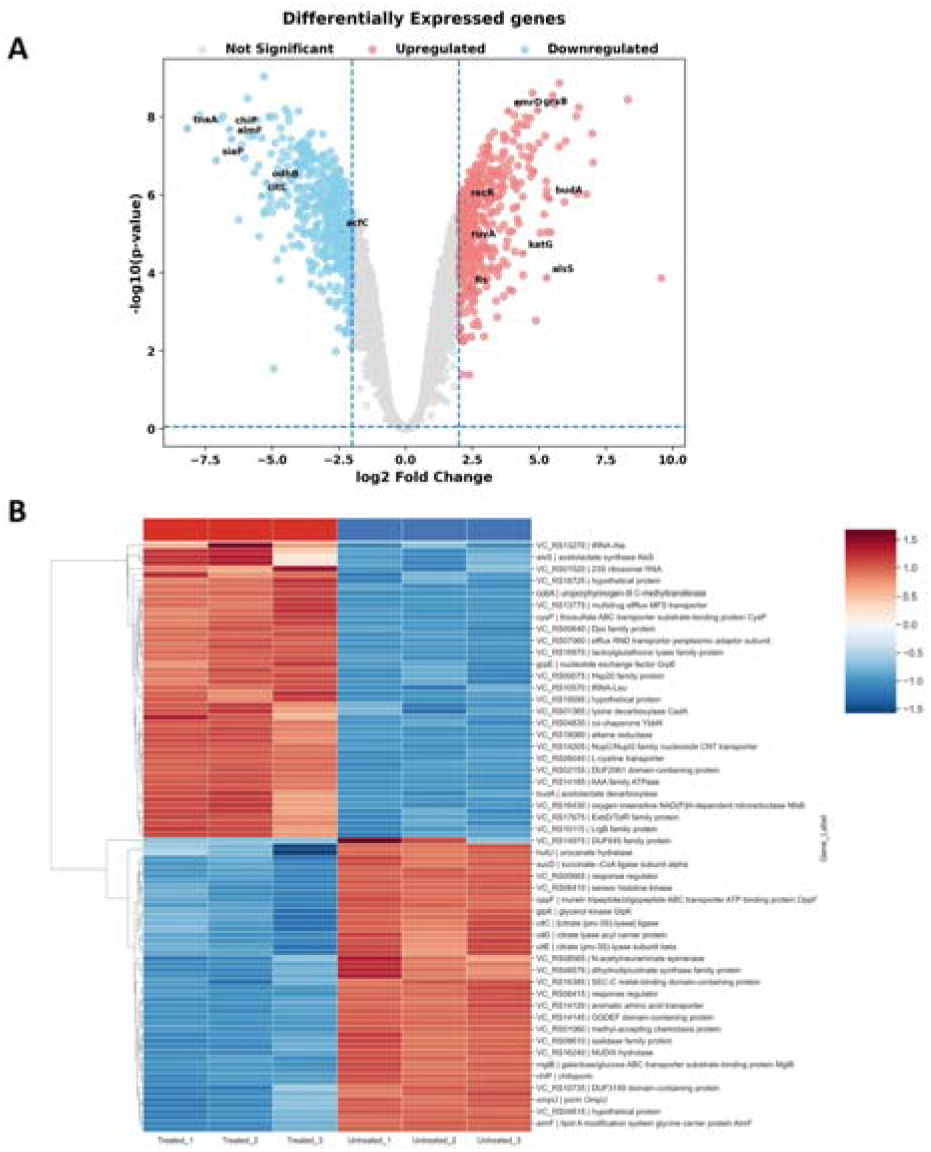
Volcano plot and hierarchical clustering heatmap of differentially expressed genes (DEGs) following L-AA treatment: **(A)** Volcano plot of L-AA-treated vs untreated samples showing significantly upregulated (red) and downregulated (blue) differentially expressed genes (DEGs), while non-significant genes are shown in grey. **(B)** Hierarchical clustering heatmap of DEGs illustrating distinct expression patterns between L-AA-treated and untreated samples, where red and blue indicate relatively high and low expression values, respectively

The heatmap represents the top 50 DEGs in *V. cholerae* following L-AA treatment compared to the untreated control. A clear separation between treated and untreated samples was observed, indicating distinct transcriptional profiles. Several genes associated with stress response, cellular adaptation, metabolic processes, and transport systems, including *alsS, budA, grpE, cysP*, and *cobA*, were upregulated in the treated group. In contrast, genes related to transport, chemotaxis, citrate metabolism, and regulatory functions, such as *oppF, citC, citD, mlpB*, and *ompU*, were downregulated in treated samples compared to untreated controls (**Figure 2B)**.

Gene Ontology (GO) was performed to analyse biological functions using p-values <0.05. The top 15 most enriched genes were identified. We found that the gene involved in energy production, transcriptional and translation machinery and metabolic and amino acid metabolism processes were significantly affected in the biological process (BP) category. In the cellular component (CC), the genes associated with central metabolic pathways, the respiratory chain complex, and signal transduction were affected. In the case of molecular function (MF), several functions related to metabolic and energy generation enzymes, transmembrane transporter activity, and enzymatic activity were significantly impacted (**Figure 3A, B and C)**.

**Figure 3.**
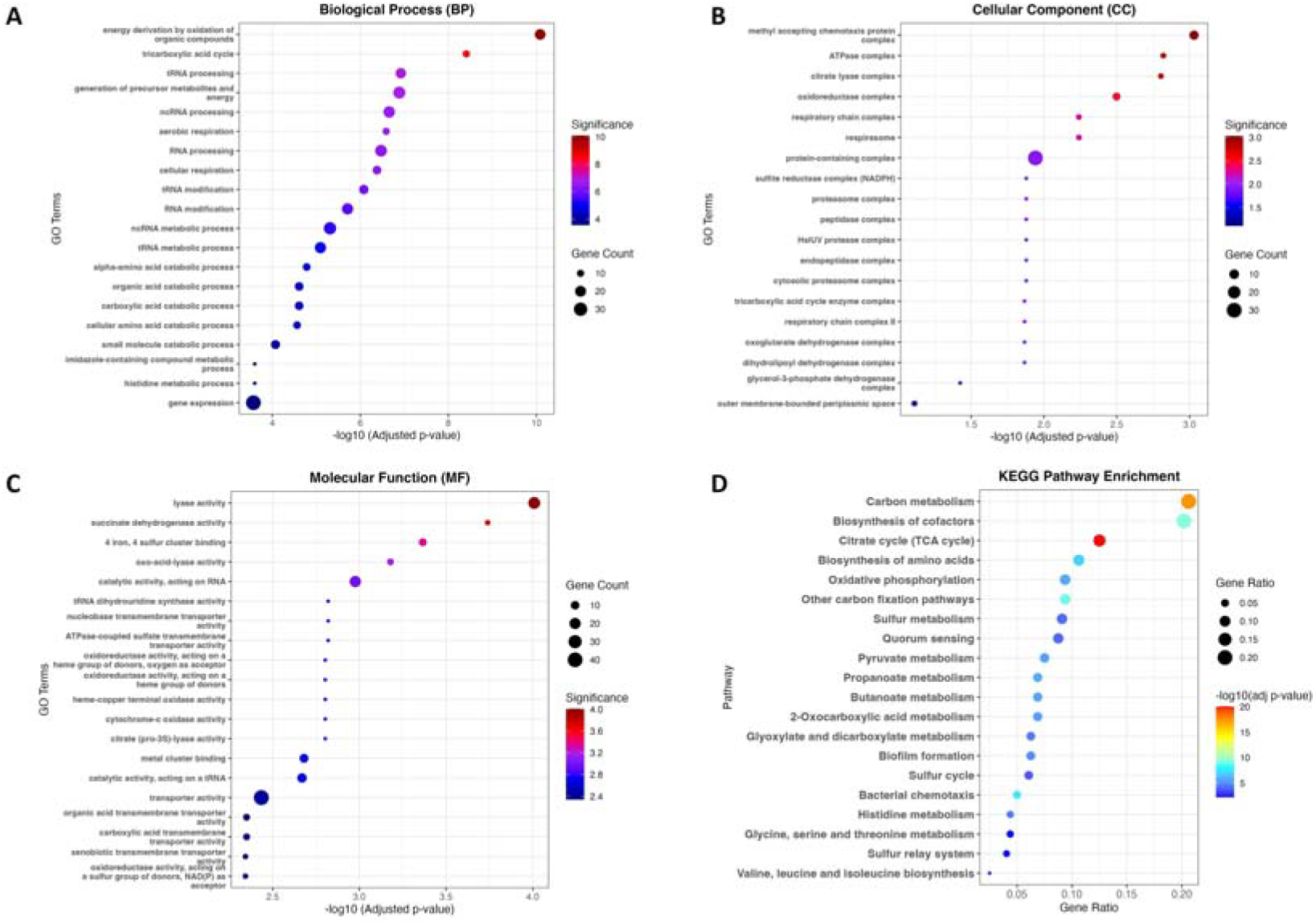
Scattered plot of GO and KEGG enrichment of Differentially expressed genes (DEGs): **(A-C)** The scatter plots of the top 15 significantly enriched Gene Ontology (GO) terms for common DEGs in terms of Biological Process (BP), cellular component (CC), and molecular function (MF) categories are shown. **(D)** The scatter plot shows the top 20 significantly enriched KEGG (Kyoto Encyclopedia of Genes and Genomes) pathways for 259 differentially expressed genes.

Out of 460 downregulated genes, KO IDs were obtained for 248 genes, of which 160 were successfully mapped to KEGG pathways. Similarly, out of 429 upregulated genes, KO IDs were obtained for 260 genes, of which 99 were mapped to KEGG pathways. KEGG pathway enrichment of the top 20 analyses revealed that differentially expressed genes were significantly expressed in metabolic pathways, particularly carbon metabolism, biosynthesis of cofactors, and the TCA cycle, indicating major disruption of energy metabolism. Additionally, pathways related to oxidative phosphorylation, sulphur metabolism, and amino acid biosynthesis were significantly affected, suggesting induction of cellular stress responses. Notably, pathways associated with quorum sensing, biofilm formation, and chemotaxis were also enriched, indicating impairment of virulence and adaptive functions in *V. cholerae* upon L-AA treatment **(Figure 3D)**. Collectively, these findings suggest that L-AA induces a global stress in *V. cholerae*, leading to activation of stress adaptation, disrupting metabolic and physiological processes, consequently compromising bacterial survival and leading to cell death.

To validate the RNA-seq data, we measured the expression levels of five representative genes (*hapR, prtV, alsS, budA* and *katG*) associated with quorum sensing, metabolism and oxidative stress with qRT-PCR. The expression levels as measured with qRT-PCR were highly in congruence with the RNA-seq data (**Figure 4**), further validating the reliability of the sequencing-based analyses.

**Figure 4.**
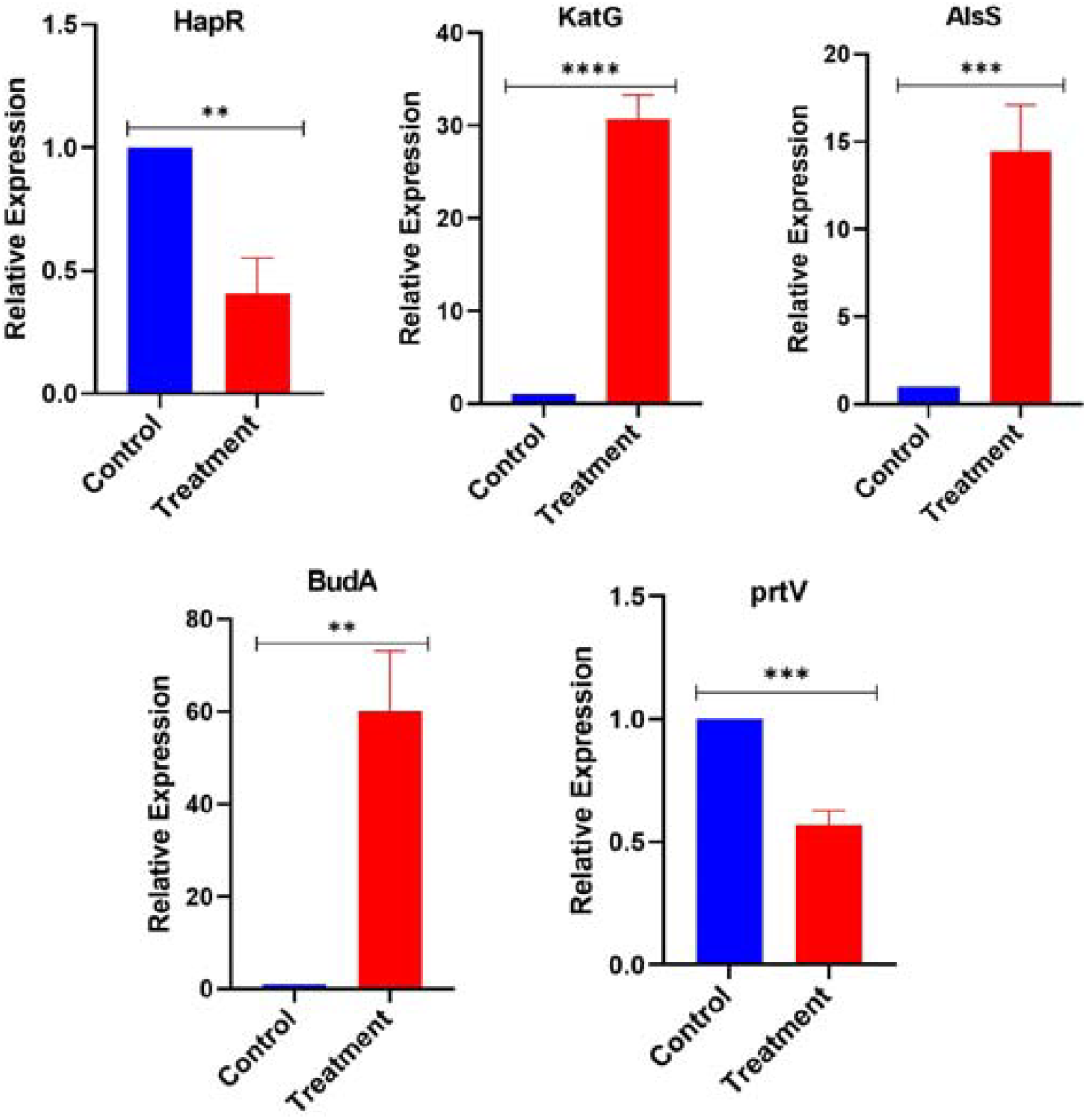
The validation of transcriptomic data by quantitative real-time PCR (qRT-PCR): The expression profile of five differentially regulated genes (*hapR, katG, alsS, budA* and *prtV*) were examined in untreated (Control) and 2.5mg/mL L-AA-treated *V. cholerae* N16961 cells. Results are presented as the mean ± SD from the three independent biological experiments. Asterisks represent significant differences as compared to control or as indicated (**** indicates P < 0.0001, *** indicates P < 0.001, ** indicates P < 0.01).

## Discussion

The anti-virulence activity of L-ascorbic acid is increasingly recognised [26]. Interestingly, vitamin C showed synergistic effects with many antibiotics, as evidenced in the UTI animal model [32]. Therefore, the antimicrobial potential of vitamin C should be harnessed therapeutically to reduce the global burden of AMR. The antimicrobial property of L-AA is linked to its prooxidant activity at high concentrations; there remains a possibility of functional modulation of gut community structure. A few interesting studies have shown its positive effect on gut community structure [33,34].

In our previous *in vitro* and *in vivo* infection model, we have shown that L-AA effectively suppresses the growth and colonization of *V. cholerae* [14,15]. The antimicrobial activity of L-AA has been well described. This study aimed to investigate the molecular basis of L-AA mediated antimicrobial activity against *V. cholerae*. Our finding demonstrates that L-AA has potent antibacterial activity by inducing oxidative and acid stress, affecting membrane integrity, and attenuating virulence-associated machinery. Several research groups have shown that excessive intracellular ROS accumulation triggers oxidative damage to lipids, proteins and nucleic acids, which leads to death in pathogens such as *Staphylococcus aureus*, carbapenem-resistant hypervirulent *Klebsiella, Mycobacterium tuberculosis, Salmonella* spp. and *V. fluvialis* [26-28,35]. The increased intracellular ROS production and compromised membrane integrity observed in the present study clearly show that oxidative stress contributes to the antibacterial effect of L-AA against *V. cholerae* **(Figure 1)**. Consistent with these phenotypic changes, transcriptomic analysis further reveals that L-AA induced a global stress response in *V. cholerae* by activating multiple stress pathways. Upregulation of *katG* and *grxB*, which suggests activation of the antioxidant defence, while induction of *cadB, alsS*, and *budA* indicates activation of acid tolerance and acetoin biosynthesis pathways that help the bacterium to maintain the intracellular pH under acidic conditions [36,37]. Upregulation of *recR, ruvA*, and the efflux transporter *emrD* further indicates activation of DNA repair and cellular protection mechanisms in response to L-AA-induced stress. Despite these adaptive responses, the extensive structural damage observed indicates that these defence mechanisms are insufficient to restore cellular homeostasis.

A key finding of this study is the coordinated repression of several virulence-associated genes. The downregulation of the quorum-sensing regulator *hapR*, along with *cqsA* and *cqsS*, suggests that L-AA may interfere with bacterial communication. HapR is the master regulator of quorum sensing, which regulates numerous genes involved in virulence, Biofilm, bacterial dispersal by producing protease and environmental adaptation [38,39]. Likewise, reduced expression of *prtV* and *hlyA*, which encode the metalloprotease and haemolysin, respectively, the reduced expression of these genes suggests diminished capacity for cytotoxicity and host tissue damage [40,41]. Furthermore, *vasJ, an* essential structural component of the type VI secretion system (T6SS), suggests that L-AA may impair bacterial competition and host interaction. Likewise, the downregulation of *acfC* encoding periplasmic accessory colonizing factor C indicates a compromised ability of *V. cholerae* to colonise the host [42,43]. The GO and KEGG enrichment analyses revealed significant alterations in energy metabolism, carbon metabolism, oxidative phosphorylation, amino acid biosynthesis, quorum sensing, biofilm formation, and chemotaxis. Collectively, these findings underscored that L-AA induces extensive metabolic shifts, forcing bacterial cells to divert resources toward stress adaptation while compromising growth and virulence **(Figures 2 and 3)**. Taken together, our results demonstrated that L-AA modulates the overall physiology and pathogenic potential of *V. cholerae*.

The global burden of cholera has increased to a concerning level, according to the WHO [44]. There is an urgent need to develop effective strategies for managing the disease. In this context, L-ascorbic acid has demonstrated its effectiveness in controlling the illness. Future research should focus on evaluating its efficacy in suitable animal model systems. In addition to managing cholera infection, the impact of L-ascorbic acid in the overall gut microbiome should be assessed. Additional studies are necessary to address these issues.

## Authors’ contribution

SRC conceived the idea and designed experiments with MK and HS. MK carried out biochemical, microscopic and RNA-related experiments. SP performed the ROS assay. MK, HS, SB and SRC analysed the data. SG analysed transcriptomic data. SRC and MK wrote the manuscript. All authors gave editorial input and approved the final manuscript.

## Funding

This work was supported by grants from the in-house (OLP-193). Manpreet Kaur, Sakshi Gautam, Suman Paul, and Himanshu Sen acknowledge CSIR for their fellowships.

## Conflict of Interest

The authors declare no conflict of interest

## Data availability statement

The data generated for transcriptomics analyses can be accessed through the NCBI Sequence Read Archive (SRA) database using reference ID PRJNA1461801. Microbiological and biochemical assay data are available on request with Dr. Saumya Raychaudhuri.

## Acknowledgements

We gratefully acknowledge Prof. Andrew Camelli, Tufts University, for generously providing the strain.

## Notes

### Competing Interest Statement

The authors have declared no competing interest.

